# The prophylactic value of TNF-α inhibitors against retinal cell apoptosis and optic nerve axon loss after corneal surgery or trauma

**DOI:** 10.1101/2022.10.06.510713

**Authors:** Eleftherios I. Paschalis, Chengxin Zhou, Jyoti Sharma, Thomas H. Dohlman, Sarah Kim, Fengyang Lei, James Chodosh, Demetrios Vavvas, Arto Urtti, George Papaliodis, Claes H. Dohlman

## Abstract

**Background and Purpose:** Late secondary glaucoma is an often-severe complication after acute events like anterior segment surgery, trauma, infection, etc. TNF-α is a major mediator that is rapidly upregulated, diffusing also to the retina and causes apoptosis of the ganglion cells and degeneration of their optic nerve axons (mediating steps to glaucomatous damage). Anti-TNF-α antibodies are in animals very effective in protecting the retinal cells and the optic nerve—and might therefore be useful prophylactically against secondary glaucoma in future such patients.Here we evaluate 1) *<u>toxicity</u>* and 2) *<u>efficacy</u>* of two TNF-α inhibitors (adalimumab and infliximab), in rabbits by *<u>subconjunctival</u>* administration.

**Methods:** For drug *<u>toxicity</u>*, animals with *<u>normal, unburned</u>* corneas were injected with adalimumab (0.4, 4, or 40 mg), or infliximab (1, 10, or 100 mg). For drug *<u>efficacy</u>*, other animals were subjected to alkali burn before such injection, or steroids (for control). The rabbits were evaluated clinically with slit lamp and photography, electroretinography, optical coherence tomography, and intraocular pressure manometry. A sub-set of eyes were stained *ex vivo* after 3 days for retinal cell apoptosis (TUNEL). In other experiments the optic nerves were evaluated by paraphenylenediamine staining after 50 or 90 days. Loss of retinal cells and optic nerve degeneration were quantified.

**Results:** Subconjunctival administration of 0.4 mg or 4.0 mg adalimumab were well tolerated, whereas 40.0 mg was toxic to the retina. 1, 10, or 100 mg infliximab were also well tolerated.Analysis of the optic nerve axons after 50 days confirmed the safety of 4.0 mg adalimumab and of 100 mg infliximab.

For *<u>efficacy, 4.0 mg adalimumab subconjunctivally in 0.08 mL</u>* provided practically full protection against retinal cell apoptosis 3 days following alkali burn, and infliximab 100 mg only slightly less. At 90 days following burn injury, control optic nerves showed about 50% axon loss as compared to 8% in the adalimumab treatment group.

**Conclusions:** *<u>Subconjunctival injection of 4.0 mg adalimumab</u>* in rabbits shows no eye toxicity and provides excellent neuroprotection, both short (3 days) and long-term (90 days). *<u>Ourtotal accumulated data from several of our studies, combined with the present paper, suggest that corneal injuries, including surgery, might benefit from routine administration of anti-TNF-α biologics to reduce inflammation and future secondary glaucoma</u>*.

## Introduction

It is well known and well documented by clinical ophthalmologists that acute traumatic events to the cornea (e.g. chemical burns, ruptured globes, infections), as well as standard penetrating corneal surgery (such as transplantations, keratoprosthesis (KPro), lacerations, etc.), often develop a late sight-threatening optic nerve neuropathy that phenotypically appears similar or identical to chronic open-angle or closed-angle glaucomas (Becker & Shaffer, 1961; Irvine & Kaufman 1969; Thoft et al., 1974; Foulks 1987; Netland et al., 1998; Quigley 1999; Aldave et al., 2000; Ayyala 2000; Kuckelkorn et al., 2001; Yaghouti et al., 2001; Rumelt et al., 2002; Quigley 2006; Tsai et al., 2006; Harissi-Dagher & Dohlman 2008; Kumar et al., 2009; Cade et al., 2011; Al-Mahmood et al., 2012; Kamyar et al., 2012; Lin et al., 2012; Talajic et al., 2012; Crnej et al., 2014; Haddadin & Chodosh 2014; Weinreb et al., 2014; Iyer et al., 2015; Wu & Xu 2017; Ali et al., 2018; Baltaziak et al., 2018; Dohlman et al., 2020; Liesenborghs et al., 2020; Geoffrion & Harissi-Dagher 2021). However, due to having an identifiable triggering cause, this category has often been labeled “secondary glaucoma.” Based on our clinical observations, secondary glaucoma may be the most consequential complication after corneal surgery.

The magnitude of this secondary glaucomatous complication has most likely been underestimated in the past due to its frequently delayed manifestation, sometimes many years (“time bomb”). Epidemiological studies on secondary glaucoma are therefore limited and numbers citing incidence and outcomes vary substantially with geography and level of economy of the area (Doren et al., 1990; Thylefors & Négrel 1994; Dana et al., 1995; Suleiman et al., 2004; Dubey et al., 2019; Gong et al., 2021; Li et al., 2021;). One source estimated that about 6 million patients in the world have secondary glaucoma compared with 67 million with the primary glaucoma (Quigley & Broman 2006). In totality, the prevalence of secondary glaucoma across the world has been stated to vary from 6 to 22% among various glaucoma studies (Dubey et al., 2019). The WHO has estimated the prevalence of *<u>blindness</u>* from secondary glaucoma to be 2.7 million people worldwide (Thylefors & Négrel 1994). Glaucomatous blindness is of course presently irreversible.

The immediate cause of secondary glaucoma, according to many studies, has been primarily attributed to surgery or trauma with corresponding inflammation rather than to chronic diseases (Gong et al., 2021)—pointing to “an acute single-event episode.” Such an acute episode may require only a relatively short period of treatment, including prophylaxis against complications such as secondary glaucoma. The majority of postsurgical glaucoma has been described as unilateral, but the eventual visual outcome can be severe (Dubey et al., 2019).

With regards to the pathophysiology of secondary glaucoma there have been recent shifts of view. Elevated IOP was almost universally blamed in the past, especially in cases of angle closure with markedly elevated pressure. However, difficulties in explaining glaucomatous damage fully in the presence of “normal” pressure has led to a greater interest in neuroinflammation and genetics (Quigley et al., 1989; Harwerth et al., 2004; Dohlman et al., 2009; Wiggs 2015; Williams et al., 2017; Dohlman et al., 2018; Wei et al., 2019; Mélik et al., 2020). In fact, an intensive research effort has been directed towards key inflammatory mediators for glaucoma in general, rather than strictly mechanical (IOP) factors (see a review)(Williams et al., 2017). The TNF-α pathway has received attention in its involvement in these processes.

Our observations in the 1990ies of the often-dramatic effect of anti-TNF-α antibodies in preventing destructive cornea melt around keratoprostheses in autoimmune patients (Dohlman et al., 2002, 2009, 2018, 2022 Ciralsky et al., 2010; Robert et al., 2017) stimulated a series of experimental studies on the pathophysiology of such treatment (Cade et al., 2014; Crnej et al., 2016; Paschalis et al., 2017, 2019; Dohlman 2022; Zhou et al. 2023). With secondary glaucoma in mind, we focused on the model of alkali burns of the cornea. These studies showed that an alkali burn can upregulate TNF-α anteriorly which will very rapidly diffuse to the retina and result in considerable retinal cell death, and degeneration of the nerve axons (the “hallmarks of glaucoma”). It had earlier been shown in studies by Kinoshita et al. that other cytokines could reach the retina in early stages after corneal burn (Miyamoto et al., 1998). (The alkali itself cannot reach the retina—it is effectively buffered at the iris plane (Kompa et al., 2005; Cade et al., 2014; Paschalis et al., 2017). These events occur very rapidly while IOP is still normal or low, pointing to the existence of a rapid, inflammatory, IOP-independent pathway to secondary glaucomatous damage after acute events elsewhere in the eye (Dohlman et al., 2019). These results have later been corroborated elsewhere (Huang et al., 2022).

Thus, at present, it is not known clinically how much of any late secondary glaucoma may be due to this newly identified potential pathway in contrast to the classic IOP-dependent influence, but it would be advisable to probe further, especially since *<u>prophylactic prevention</u>* should be a possibility since drugs are already available and can be promptly applied. Thus, importantly, it has been shown that not only corticosteroids but also monoclonal antibodies (mAbs) to TNF-α, such as etanercept, infliximab, or adalimumab, can be markedly neuroprotective to the retina in animal injury models if administered rapidly enough systemically after the instigating event (Roh et al., 2012; Cade et al., 2014; Paschalis et al., 2017). In recently published animal work, we have further demonstrated that prolonged administration of anti-TNF-α antibody to the retina can be achieved by subconjunctival implantation of a polymer-based drug delivery system (DDS) (Robert et al., 2016; Zhou et al., 2023). Although the DDS was loaded with only 85 μg of infliximab (∼30 μg/kg), a biological effect was observed for over a month, manifested by a significant reduction in retinal cell apoptosis after an alkali burn to the cornea (Zhou et al., 2017).

However, of considerable importance are also previous findings in animals that not only the described rapid inflammatory response can be suppressed by the biologics but also, quite likely, the classic IOP-triggered insult as well, since the optic nerve degeneration found after 3 months could have resulted from either or both pathways. Several of these studies showed a very marked effect of the antibodies to protect the retinal cells and the optic nerve (50-100% protection), strengthening the possibility of prophylaxis of secondary glaucoma at the time of the inflammatory event (surgery, trauma, etc.), and perhaps also shortly afterwards. These findings alone should have substantial therapeutic promise.

Because of these insights and therapeutic possibilities, it seems that further investigations of the ocular use of anti-inflammatory biologics are warranted. Not only postoperative glaucoma is at stake but also other inflammatory complications such as corneal tissue melt, retroprosthetic membranes, uveitis, vitritis, retinal detachments, etc. This study is meant to focus specifically on the following issues: The risks and benefits of the two presently leading marketed TNF-α inhibitors (adalimumab and infliximab), when administered locally to the eyes, should be clarified. Also, it is still not clear whether a drug delivery system (DDS) is the most practical way for the administration, or whether a single injection of an agent would suffice—and be safer and more practical—or whether an intravitreal or systemic route would still be preferable for long-term use. The present study has therefore attempted to investigate the toxicity and efficacy of such anti-inflammatory administration by the *<u>subconjunctival</u>* route.

## Materials and Methods

### Rabbit model

All animal-based procedures were performed in accordance with the Association for Research in Vision and Ophthalmology Statement for the Use of Animals in Ophthalmic and Vision Research, and the National Institutes of Health Guidance for the Care and Use of Laboratory Animals. This study was approved by the Animal Care Committee of the Massachusetts Eye and Ear Infirmary and Schepens Eye Research Institute. Dutch-belted pigmented rabbits were used for this study and were obtained from Covance (Dedham, MA, USA). Rabbits were used at the ages of 4-10 months. For safety studies, animals were divided into the following groups, saline group (n=1), subconjunctival administration of 100 mg infliximab (n=1), triamcinolone (n=3), 4 mg adalimumab (n=3) and 40 mg adalimumab (n=2). For the pilot efficacy studies, saline group (n=1), 100 mg infliximab (n=1), 20 mg triamcinolone (n=3), 4 mg adalimumab (n=3) and 40 mg adalimumab (n=2). For long-term efficacy studies, saline group (n=3) and 4 mg adalimumab (n=3).

### Rabbit anesthesia, recovery, and euthanasia

Rabbits were anesthetized with intramuscular injection of ketamine hydrochloride INJ, USP (35 mg/kg; KetaVed, VEDCO, St. Joseph, MO, USA), xylazine (5 mg/ kg; AnaSed, LLOYD, Shenandoah, IA, USA), and acepromazine (0.75 mg/kg; PromAce®, Boehringer Ingelheim Vetmedica, Inc. MO, USA). Reversal of anesthesia was obtained with intravenous (IV) yohimbine (0.1 mg/kg; Yobine, LLOYD) administration in a marginal ear vein. Rabbits were placed on a warm pad until becoming sternal and able to move. Anesthetized rabbits were euthanized at the completion of the experiment with 100 mg/kg Fatal Plus IV injection (sodium pentobarbital).

### Retinal injury

Sterile inflammatory retinal injury was accomplished by applying a corneal surface alkali burn, an established retinal injury model (Cade et al., 2014; Paschalis et al., 2017). Topical anesthetic (0.5% proparacaine hydrochloride, Bausch & Lomb, Tampa, FL, USA) was applied to the study eye while the contralateral eye was protected using GenTeal gel (Alcon, Fort Worth, TX, USA). Alkali burn was performed by using an 8-mm diameter filter paper soaked in 2 N NaOH that was applied to the center of the cornea for 10 seconds followed by immediate eye irrigation with saline solution for 15 minutes. Buprenorphine (0.03 mg/kg; Buprenex Injectable, Reckitt Benckiser Health- care Ltd, United Kingdom) was administered subcutaneously prior to the burn procedure for pain management and a transdermal fentanyl patch (12 mcg/hr; LTS Lohmann Therapy System, Corp., NJ, USA) was placed on the right skin to alleviate pain for 3 days.

### Dose titration of anti-TNF-α and monoclonal antibodies

Two FDA approved TNF-α inhibitors were selected: a) infliximab (Remicade® Janssen Biotech Inc., Johnson and Johnson, Titusville, NJ, USA), and b) adalimumab (Humira® Abbvie Inc., North Chicago, IL, U.S.A.). Infliximab was administered subconjunctivally at 1 mg (0.008 mL), 10 mg (0.08 mL), or 100 mg (0.8 ml) doses (n=3) in otherwise intact eyes immediately after the irrigation of alkali injury. Likewise, adalimumab was administered subconjunctivally at 0.4 mg (0.008 mL), 4 mg (0.08 mL), and 40 mg (0.8 mL) doses (n=3). Two sham injection (control) animals were used (one for each study group) that received 0.8 mL of sterile saline subconjunctivally without drug. Subconjunctival injections of infliximab were performed using a 30-G needle and adalimumab using the pre-fitted syringe needle.

### Clinical Evaluation

Clinical evaluation was performed on all rabbits before and after treatment and 0.5% proparacaine hydrochloride was applied to the operated eyes. Eyes were photographed using a digital SLR camera (Nikon, Tokyo, Japan) attached to a surgical microscope (S21; Carl Zeiss, Jena, Germany). Remote photography was performed using an iPhone 7plus (Apple Inc) fitted with a magnifying clip-on lens (12x, Pictek Fisheye Lens, Amazon.com Inc, WA, USA).

### IOP measurements

Intraocular pressure measurements were performed in anesthetized rabbits using a custom-made intracameral pressure transducer connected to a 27-gauge needle. The device was designed using a differential microelectromechanical pressure sensor 40 PC (Honeywell, Freeport, IL) connected to a 14-bit, 48 kilo samples per second data acquisition NI USB-6009 (National Instruments, Austin, TX), controlled by a proprietary software algorithm operating in Labview 2017 (National Instruments) environment. A special algorithm was designed to compensate for aqueous humor volume displacement during *in vivo* pressure measurements. The device was assembled using microfluidic components (IDEX Health & Science, Oak Harbor, WA) with minimum dead volume. Before measurements, the remaining dead volume of the syringe was pre-filled with sterile water, thus minimizing air compressibility only within the micro- electromechanical cavity, which was co-evaluated by the software algorithm. To perform measurements, the needle was inserted into the anterior chamber of the eye through a temporal clear corneal puncture, adjacent to the limbus, and the needle was advanced approximately 5 mm toward the center of the chamber (Paschalis et al., 2019).

### Electroretinography

Animals were dark-adapted in a ventilated dark chamber for 30 minutes, followed by anesthesia. The rabbits were placed on a warm platform (38°C) and both eyes were dilated using Tropicamide Ophthalmic Solution USP, 1% followed by Proparacaine Hydrochloride Ophthalmic Solution USP, 0.5% topical drops. GenTeal was applied to both corneal surfaces and a contact lens electrode was fitted to the right eye. A reference needle electrode was placed into the forehead above the midline of the eye and a ground needle electrode above the rabbit’s tail.

Dark-adapted ERGs were performed as follows:

Step 1: 6 sweeps, 5000 ms inter sweep delay, 1 Hz frequency, 0.01 cd.s/m2 intensity on for 4 ms of color White-6500K.

Step 2: 6 sweeps, 15000 ms inter sweep delay, 1 Hz frequency, 3 cd.s/m2 intensity on for 4 ms of color White-6500K.

Step 3: 6 sweeps, 20000 ms inter sweep delay, 1 Hz frequency, 10 cd.s/m2 intensity on for 4 ms of color White-6500K.

After completion of dark-adapted ERG in both eyes, the animals were light-adapted in ambient room light for 10 minutes, followed by light-adapted ERG using the following steps:

Step 1: 10 sweeps, 2000 ms inter sweep delay, 1 Hz frequency, 10 cd.s/m2 intensity on for 4 ms of color White-6500K with a background light of 30 cd/m2 of color White-6500K

Step 2: 50 sweeps, 30 ms, 30 Hz frequency, 10 cd.s/m2 intensity on for 4 ms of color White- 6500K with a background light of 20 cd/m2 of color White-6500K ERG measurements were performed using the E3 Electrophysiology System (Diagnosys LLC, MA, USA) with full-field binocular desktop Ganzfeld and analyzed using the Epsilon V6 (Diagnosys LLC, MA, USA) software. A-wave amplitude was measured as the orthogonal distance of the negative dip from baseline (reference). b-wave amplitude was measured as the absolute orthogonal distance of the a from the first elevation. Time response was measured for the light stimuli. Similar protocol was used for light-adapted ERG measurements.

### *In vivo* optical coherence tomography

Posterior segment optical coherent tomography (OCT) was performed in anesthetized animals using the Heidelberg Spectralis (Heidelberg Engineering GmbH, Heidelberg, Germany). A speculum was used to retract the lids. Vertical and horizontal raster scans were performed to acquire images of the 4 retinal quadrants adjacent to the optic nerve (ON). Radial scans were performed to image the ON. Quantification of retinal thickness was performed with image segmentation using ImageJ software version 1.43 or above (NIH, Bethesda, MD; http://imagej.nih.gov/ij). Thickness was measured from the borders of the ganglion cell layer to the outer nuclear layer. Six images of superior, inferior, temporal, and nasal retina were measured. Baseline images were compared to images obtained 50 days after subconjunctival injection.

### Tissue preparation

Eyes were enucleated at predetermined time points. The eyes were surgically dissected and fixed in 4% paraformaldehyde (PFA) (Sigma-Aldrich, St Louis, MO, USA) solution for 3 days at 4°C. Following fixation, eyes were sagittally dissected and half of the eye ball was embedded in optimal cutting temperature (OCT) compound and flash-frozen, and the other half in glycol methacrylate. OCT embedded tissues were sectioned at 10 μm thickness and glycol methacrylate embedded at 3 μm and transferred to positively charged glass slides (Superfrost glass slides, Thermo Fisher, IL, USA). Hematoxylin/eosin (H&E) staining was performed for general histologic observation. 10 μm cryosections were used for TUNEL assay and 3 μm methacrylate sections for H&E. Cryosections were prepared only for the analysis of retinal cell death.Histological/ morphological studies were performed in methacrylate embedded tissues to quantify retinal cell number.

### Retinal damage and cell death

Cell death was assessed in tissue sections using terminal deoxynucleotidyl transferase-mediated dUTP nick- end labeling (TUNEL, Roche TUNEL kit (12156792910; Roche, Basel, Switzerland), as previously described (Cade et al., 2014; Paschalis et al., 2017). Mounting medium with DAPI (Ultra- Cruz; sc-24941; Santa Cruz Biotechnology, Dallas, TX) was placed over the tissue, followed by a coverslip. Tile images were taken using an epifluorescent microscope (Zeiss Axio Imager M2; Zeiss, Oberkochen, Germany). DAPI signal (blue) was overlayed with Texas red (TUNEL+ cells) and quantified with ImageJ software version 1.43 or above (NIH, Bethesda, MD; http://imagej.nih.gov/ij) to assess the number of TUNEL+ cells overlapping with DAPI in the areas of interest. At least 3 different tissue sections per eye were analyzed, and data were presented as a percentage of the total DAPI area. Quantification of TUNEL+ cells was preformed centrally at 1/3^rd^ of the diameter of the retina and peripherally at 2/3rds of the diameter.

### Optic nerve evaluation with paraphenylenediamine staining

Optic nerve axon degeneration was evaluated in enucleated rabbit eyes using paraphenylenediamine staining (PPD). Optic nerves were dissected from the enucleatedeyes, fixed in Karnovsky fixative solution for 24 hours at 4°C, then processed and embedded in acrylic resin. Tissue cross sections (1 μm thick) were stained with 1% PPD in absolute methanol. Each section was mounted onto a glass slide and imaged using a bright field microscope (Nikon eclipse E800 DIC; Tokyo, Japan) with a 100X objective lens. Tile images of the whole nerve section were obtained, and axon degeneration was quantified using ImageJ software, according to previous protocols (Paschalis et al., 2017).

### Retinal neuroprotection

The efficacy of subconjunctival anti-TNF-α administration in retinal neuroprotection was assessed using the aforementioned ocular burn injury model (Cade et al., 2014; Paschalis et al., 2017). Immediately after the burn and lavage, the eye received subconjunctival injection of either infliximab (1, 10, or 100 mg) or adalimumab (0.4, 4, or 40 mg). Sterile saline (sham) subconjunctival injection (0.8 mL, n=3) was used as control. Additional controls included: subconjunctival injection of triamcinolone (20 mg, n=3). Retinal protection was assessed using TUNEL assay. Long-term efficacy of subconjunctival 4 mg adalimumab was assessed in rabbit eyes, 90 days after injury using retinal H&E and ON PPD staining. The mid periphery region of H&E-stained retina and PPD stained nerve section was imaged using bright field microscopy with a 63X objective lens. The data was analyzed by counting the cells in different retinal layers using Image J software 1.43 or above (NIH, Bethesda, MD; http://imagej.nih.gov/ij). The cells were counted manually in a defined area at different sections, averaged and normalized using the contralateral eye. Axon degeneration was quantified using ImageJ software, per previous protocols (Paschalis et al., 2017).

### Statistical analysis

Results were analyzed with the statistical package of social sciences (SPSS) Version 17.0 (Statistical Package for the Social Sciences Inc., Chicago, IL, USA). The normality of continuous variables was assessed using the Kolmogorov-Smirnov test. Quantitative variables were expressed as mean±standard deviation (SD). The Mann-Whitney test was used to assess differences between groups. All tests were two-tailed, and statistical significance was determined at p < 0.05. The independent student t-test was used to compare means between two groups, and pairwise t-test to compare changes within the same group. Analysis of variance (ANOVA) was used for comparisons of multiple groups. Alpha level correction was applied, as appropriate, for multiple comparisons.

## Results

### Retinal toxicity study

To *<u>assess toxicity</u>*, naive Dutch-Belted pigmented rabbits received subconjunctival injection of adalimumab (0.4, 4, 40 mg) or infliximab (1, 10, 100 mg) in one eye. All injections were uneventful. A conjunctival bleb was observed immediately after the injection of a higher dose of adalimumab (40 mg) and infliximab (100 mg), which resolved within a day **(Fig. 1 A-C)**. No bleb was generated following injection of 4 mg adalimumab in 0.08 mL saline **(Fig. 1 D-F)**.

**Figure 1.**
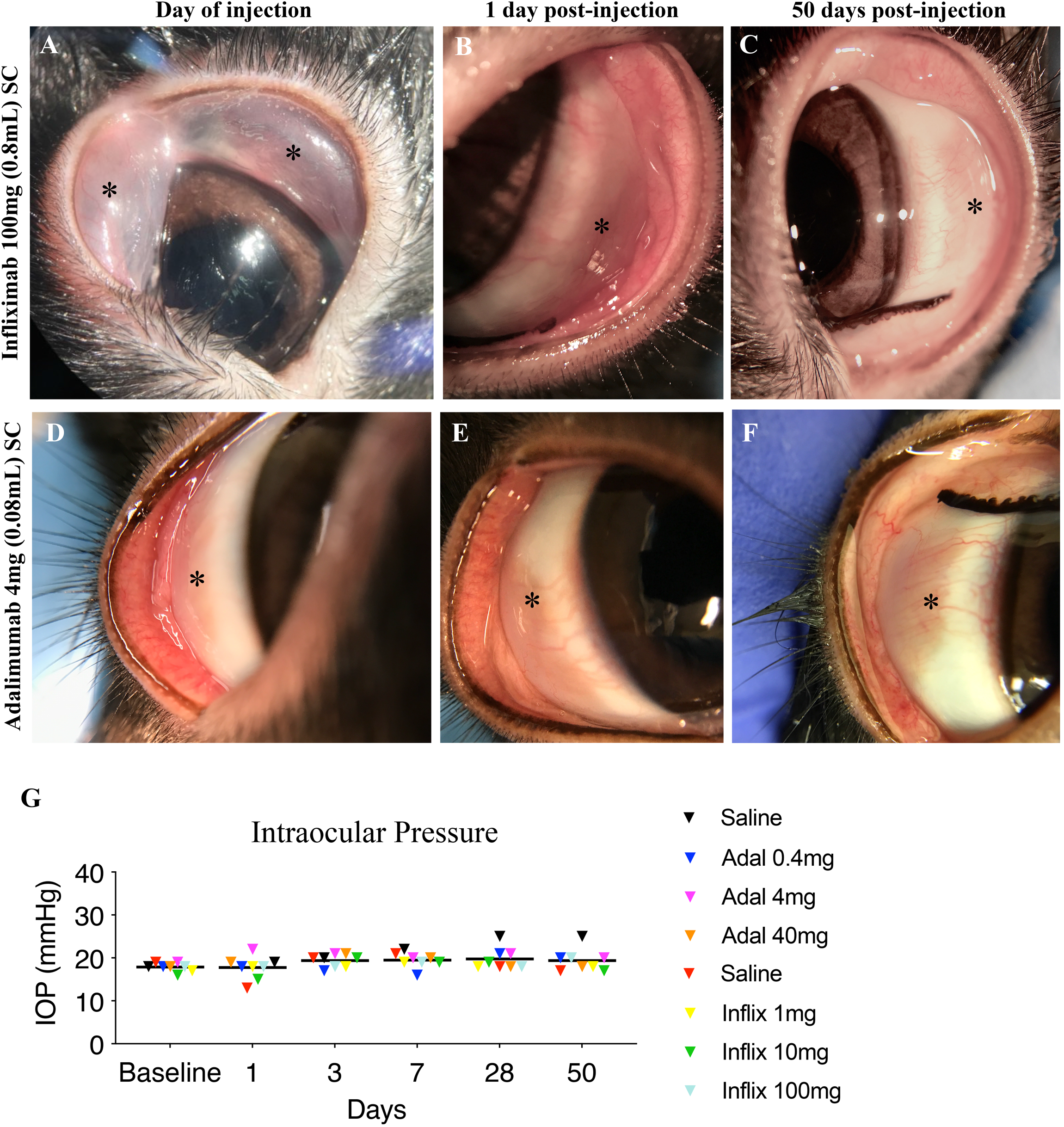
Toxicity: Subconjunctival injection of infliximab and adalimumab in normal eyes. **(A)** Photograph of subconjunctival injection of **100 mg infliximab (0.8 m)** showing elevation of the superior and inferior conjunctiva immediately after the injection as the volume was distributed equally in the superior and inferior conjunctiva. **(B)** The same eye the following day showing lack of bleb and **(C)** at 50 days with normal appearance. **(D)** Photograph of the superior subconjunctiva after injection of **4 mg adalimumab** (**0.08 ml)**. No elevation of the conjunctiva is evident immediately after the injection. **(E - F)** Normal appearance of the eye at day one and 50 days post injection. **(G)** Longitudinal intraocular pressure measurements using cannulation in eyes injected subconjunctivally with either 1, 10, or 100 mg of infliximab or 0.4, 4, or 40 mg of adalimumab. No IOP elevation is evident up to 50 days post injection. A slight elevation was observed in the saline (sham) injected eyes at day 28 and 50, which did not reach statistical significance. One animal per dose with serial measurements, total n=8.

Fifty days after injection of the agents, all eyes looked normal, with intact corneal and conjunctival epithelium and lack of signs of inflammation or vascularization **(Fig. 1 C, F)**.

Nor did subconjunctival injection of adalimumab or infliximab, at various doses, cause immediate or long-term intraocular pressure elevation, as assessed by intracameral manometry (an established and previously reported technique (Paschalis et al., 2017) 7 days before injection and 1, 3, 7, 28 and 50 days after injection **(Fig. 1 G)**. A slight IOP elevation occurred only 7, 28, and 50 days after injection of sterile saline (sham group) but was not statistically significant **(Fig. 1 G)**.

Additional evaluation was performed to assess potential retinal toxicity following subconjunctival injection of adalimumab (up to 40 mg) or infliximab (up to 100 mg). Dark-adapted and light-adapted electroretinography (ERG) were performed at baseline (7 days prior to injection) and 3, 7, 28, and 45 days post injection. Adalimumab and infliximab, at the highest doses, did not cause appreciable changes, and ERG responses following flash stimulation with 0.01, 3, or 10 cd.S/m2. **(Supplemental figures 1, 2)**.

### *In vivo* histology

*In vivo* histology was performed by using OCT imaging at baseline and 50 days after subconjunctival injection of 4 mg and 40 mg adalimumab **(Fig. 2 A, B)** and 100 mg infliximab **(Fig. 2 C)**. 4 mg of adalimumab did not cause changes in retinal thickness **(Fig. 2 D)**, while high dose (40 mg) adalimumab caused an increase in superior retinal thickness (P<0.05) as compared to baseline measurements **(Fig. 2 E)**. No retinal thickness changes were observed following subconjunctival injection of 100 mg infliximab **(Fig. 2 F)**. Lower dose adalimumab (0.4 mg) and infliximab (1 and 10 mg) likewise did not cause changes in retinal thickness measured by OCT. Representative high magnification image of superior retinal thinning after subconjunctival injection of 40 mg adalimumab **(Supplemental figures 3)**.

**Figure 2.**
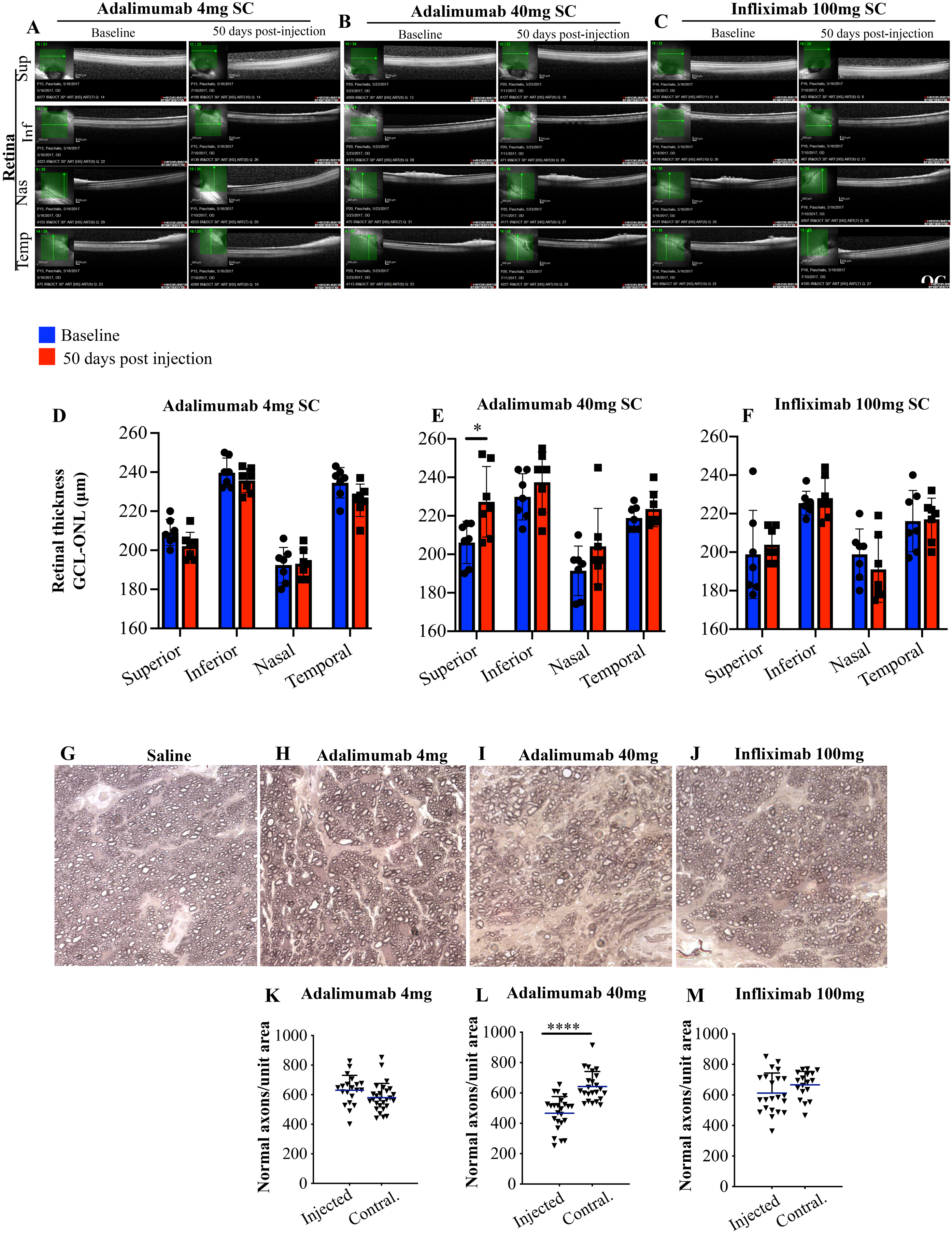
Toxicity: Retinal and optic nerve histology, 50 days after subconjunctival administration of adalimumab or infliximab in normal eyes. *In vivo* optical coherence tomography (OCT) of the 4 retinal quadrants at baseline and 50 days post subconjunctival injection with **(A)** 4 mg adalimumab **(B)** 40 mg adalimumab and **(C)** 100 mg of infliximab. Quantification of the retinal thickness from the boundaries of the ganglion cell layer (GCL) to the outer nuclear layer (ONL) shows **(D)** 4 mg adalimumab did not cause any change in the retinal thickness. **(E)** 40 mg adalimumab caused significant increase in the thickness of the superior quadrant 50 days post injection. **(F)** 100 mg infliximab did not cause any appreciable change in retinal thickness. p-Phenylenediamine (PPD) staining of the optic nerves 50 days after subconjunctival injection of **(G)** saline (sham) or **(H, I)** 4 mg adalimumab did not cause optic axon degeneration. **(J, K)** marked axonal degeneration was evident after subconjunctival injection of 40 mg of adalimumab. All comparisons were performed using the contralateral un-injected eye. **(L, M)** 100 mg of infliximab did not cause appreciable optic nerve axon degeneration. Scale bar: 50 μm. **(D-F)** Two-way ANOVA with Sidak’s correction *P<0.05, **(K-M)** Student t-test ****P<0.0001 (sections of one eye).

### Long-term toxicity study

Further toxicity assessment was performed by analyzing the optic nerve axons 50 days after subconjunctival injection of either 0.4, 4, 40 mg of adalimumab or 1, 10, 100 mg of infliximab. The results confirmed the safety of 4 mg adalimumab and the adverse effect of 40 mg adalimumab, which resulted in axonal degeneration and drop-out, as compared to the contralateral un-injected eye and control saline (sham) injected eye **(Fig. 2 D, E, G-I, K, L)**.

Infliximab 100 mg on the other hand did not cause optic nerve axonal degeneration/drop-out, consistent with the OCT retinal thickness measurements **(Fig. 2 F, J, M)**.

### Efficacy of inhibitors and neuroprotection

The efficacy of subconjunctival injection of adalimumab and infliximab in acute retinal protection was evaluated 3 days after corneal burn. We have previously shown that corneal alkali burn causes acute uveal inflammation, release of TNF-α, and subsequent retinal cell apoptosis within 3 days in rabbits (Paschalis et al., 2017). Indeed, saline treated (subconjunctival) rabbits exhibited peripheral and central retinal cell apoptosis within 3 days after the injury, extending in all 3 retinal layers **(Fig. 3 A, B, I)**. In contrast, subconjunctival injection of infliximab 100 mg reduced retinal cell apoptosis **(Fig. 3 C, D, I)** but not as effectively as subconjunctival adalimumab 4 mg, which provided complete protection to the retina **(Fig. 3 G, H, I, J)**.

**Figure 3.**
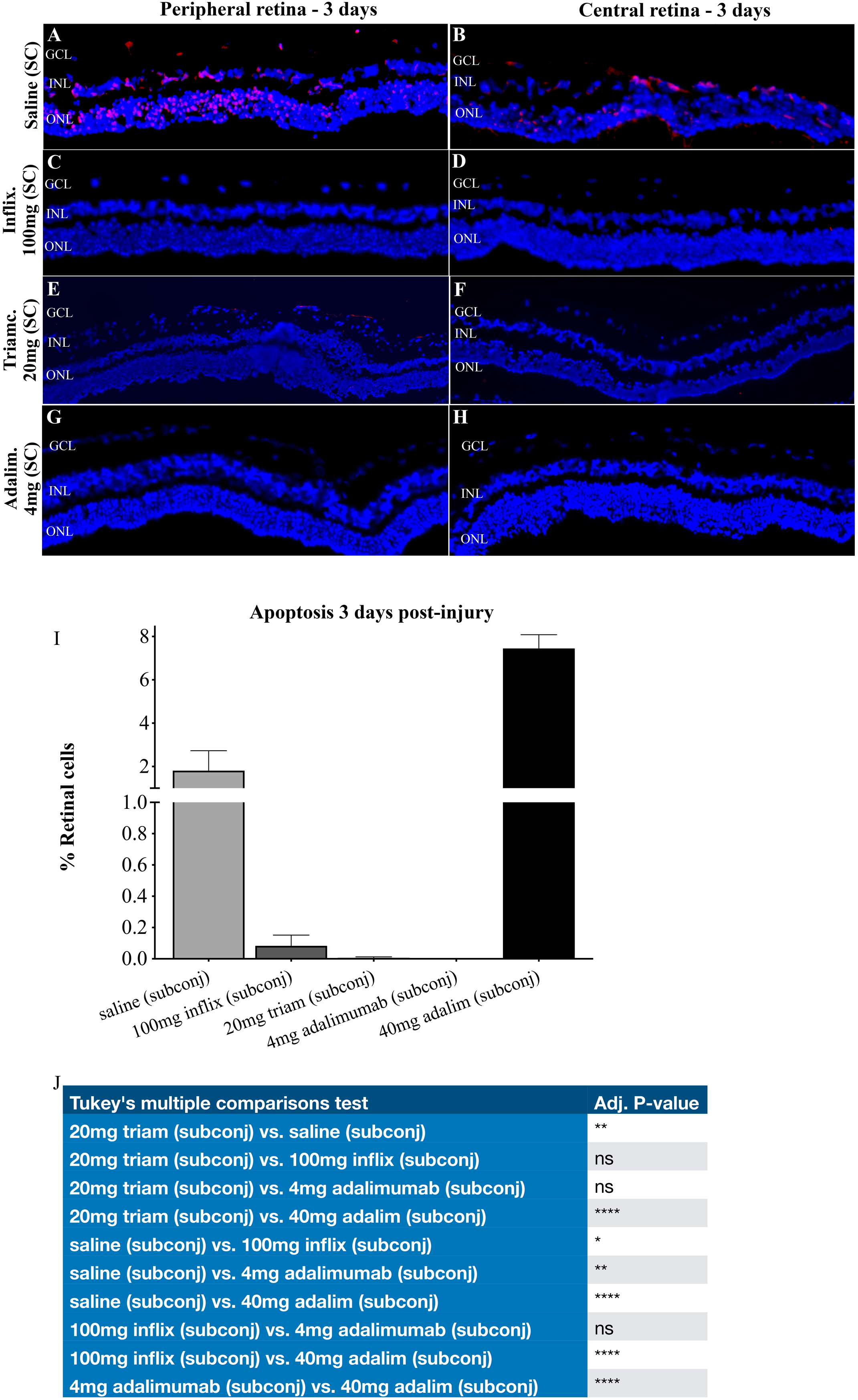
Efficacy: Pilot experiment. Acute protection (3 days) of the retina with subconjunctival adalimumab after corneal burn. Evaluation of peripheral and central retinal cell apoptosis using TUNEL assay 3 days after acute corneal surface injury with alkali. **(A, B)** Saline (sham) treatment results in significant cell apoptosis in all 3 retinal layers. **(C - D)** Subconjunctival administration of 100 mg of infliximab 15 minutes after the injury results in appreciable reduction of retinal cell apoptosis. **(E, F)** Subconjunctival injection of 20 mg triamcinolone also shows almost complete protection to the retina, with some apoptosis evident in the ganglion cell layer of the peripheral retina. **(G, H)** Subconjunctival injection of 4 mg adalimumab provides 100% retinal protection in the peripheral and central retina. **(I)** Quantification of retinal cell apoptosis following subconjunctival injection of various doses and drugs. Note that 40 mg adalimumab exacerbates cell apoptosis, indicative of its toxic effect at high dose, however, 4 mg adalimumab provides the most optimal retinal protection, as compared in all studied regimens. **(J)** Two-way ANOVA with Tukey’s correction. (n=3: 20 mg trim., 4 mg adalim; n=2: 40 mg adalim; n=1: saline, 100 mg inflix).

Likewise, subconjunctival triamcinolone 20 mg was very protective **(Fig. 3 E, F, I, J)**. In contrast, high dose (40 mg) subconjunctival adalimumab was toxic and caused retinal cell apoptosis rather than retinal protection (compared to sham control) **(Fig. 3 I, J)**.

The long-term efficacy of subconjunctival 4 mg adalimumab in retinal and optic nerve protection was evaluated in a separate study, by evaluating the retina and optic nerve 90 days after corneal burn. A single subconjunctival injection of 4 mg adalimumab after the injury was able to prevent retinal cell loss and change in retinal thickness, as compared to saline treatment, which exhibited significant loss of cells in the ganglion cell, inner nuclear, and outer nuclear layer **(Fig. 4 A- E)**. Moreover, analysis of the optic nerve axons confirmed the above results, showing that 4 mg adalimumab was able to preserve 92% of the ON axons at 90 days, as compared to saline treatment which exhibited almost 50% axonal loss **(Fig. 4 F-H)**.

**Figure 4.**
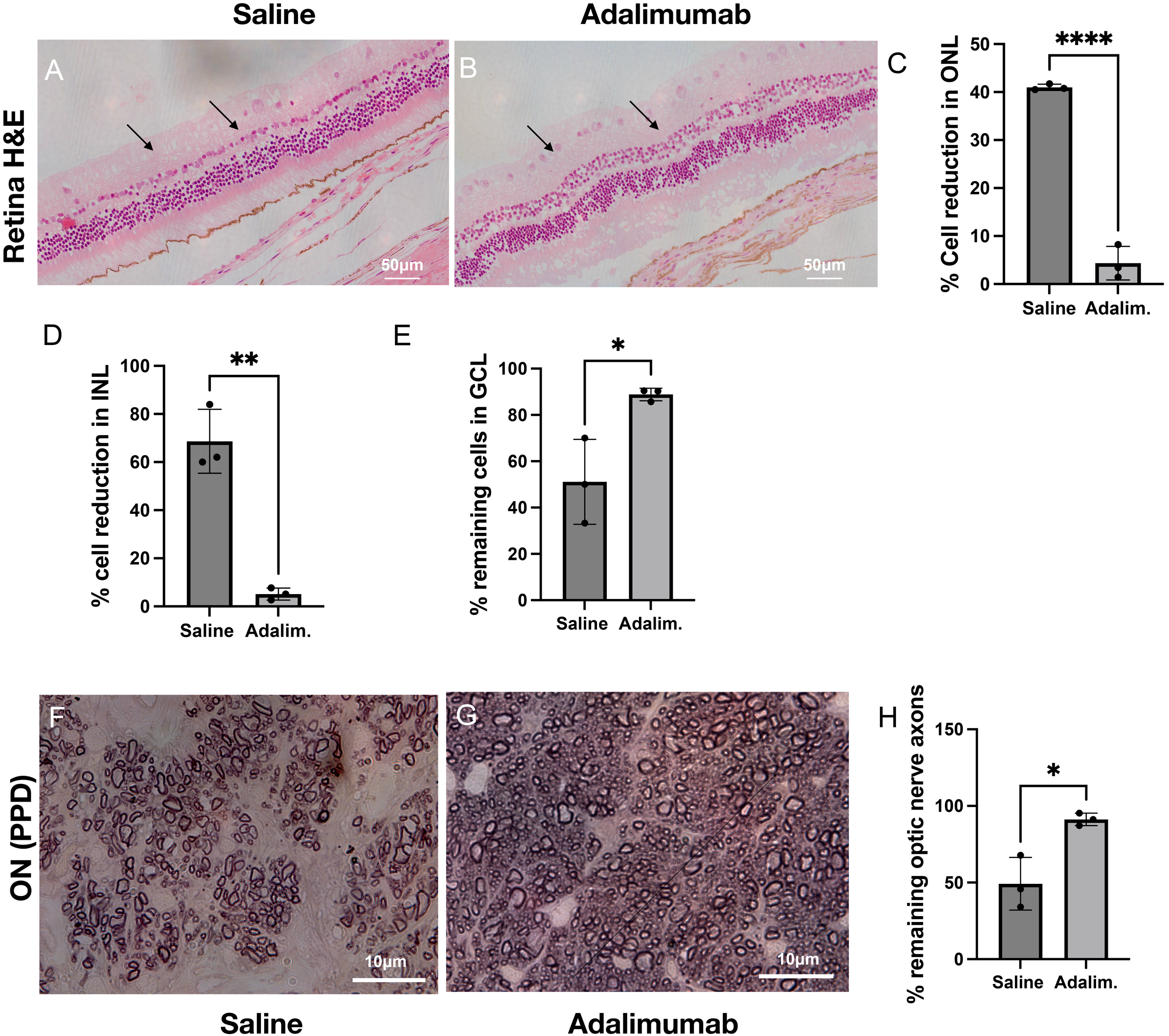
Efficacy: Pilot experiment. Long-term protection (90 days) of the retina with 4 mg subconjunctival adalimumab after corneal burn. Histologic examination (H&E) of the retinas with hematoxylin and eosin staining shows significantly increased in retinal thickness (p=0.03, n=3) and reduction in cell count in outer nuclear layer (p<0.0001, n=3), inner nuclear layer (p=0.0012, n=3) and neuronal cell count in the ganglion cell layer in the saline treated group (p=0.02, n=3), as compared to adalimumab (4 mg) treated group which exhibits minimal changes in retinal thickness and RGC count, indicative of long-term neuroprotective effect. **(A-E)** The saline (sham) treated group exhibits significant 55% reduction in optic nerve axons (p=0.014, n=3) as compared to adalimumab treated eyes that exhibit minimal 9% loss of optic nerve axons. **(F - H)** Student t-test.

### Mathematical calculations of drug bioavailability

The expected retinal bioavailability of adalimumab after subconjunctival administration has been roughly estimated computationally. It is known that about 0.06% of a hydrophilic compound (Gd-DTPA) permeates to the rabbit retina after subconjunctival injection^66^. Similar results were obtained by computational simulation of protein drug permeation from the subconjunctival space to the retina and vitreous (i.e. 0.1–1% of the dose; fraction of 0.001-0.01) (Ranta et al., 2010).

This estimate takes into account permeability in the sclera, choroid, and RPE as well as drug loss to the conjunctival and choroidal blood flows. Average retinal adalimumab concentration after subconjunctival injection can be estimated using the equation:

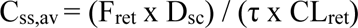

where Fret (retinal bioavailability) = 0.001-0.01, Dsc (subconjunctival dose) = 4 mg, τ (follow-up period after the dose) = 72 h (Fig. 5), and CLret (clearance from the retina/vitreous humor compartment) = 0.066 ml/h (Subrizi et al., 2019). Therefore, the concentration of adalimumab (4 mg; molecular weight 144,190 g/ml) in the retina/vitreous compartment is expected to be 0.8-8 μg/ml (5.8-58 nM); about 10^3^-10^4^ times higher than its affinity (Kd = 8.6 pM) to soluble TNF-α (Kaymakcalan et al., 2009), and 10-100 times higher than adalimumab affinity towards membrane bound TNF-α (Kd = 468 pM). Thus, it is likely that direct permeation of adalimumab from the subconjunctival space to the retina results in the observed therapeutic activity in the retina.

It is known that substantial systemic drug absorption takes place from the subconjunctival injection site. Experimental data (Kim et al., 2008) and computational follow-up analysis (Ranta et al., 2010) suggest systemic absorption of 72–83% for large molecules. Then, average steady-state concentration of adalimumab (Css,av) in human plasma and adalimumab quantity entering retina during 72 hours can be estimated as:

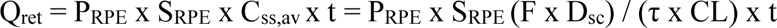

where PRPE = 0.035 x 10^-6^ cm/s (RPE permeability of bevacizumab from) (Ramsay et al., 2019), SRPE = 12 cm^2^ (area of human RPE)(Panda-Jonas et al., 1994), F = 0.72, Dsc = 4 mg, τ = 72 h, CL (plasma clearance) = 12 ml/h (Mascelli et al., 2007) and t = 72 h. Average concentration in plasma is calculated to be ≈ 3.2 mg/L (22 nM) resulting in distribution of ≈ 0.35 μg of adalimumab across the RPE into the retinal compartment within 72 hours. Permeation of proteins across the blood retinal barrier is slow (Vellonen et al., 2016) and the result (0.35 μg) is about 0.01% of the subconjunctival dose. *Therefore, we conclude that direct permeation of adalimumab across the ocular tissues to the retina represents about 10-100 higher bioavailability (≈ 0.1–1%) than its retinal entry via systemic blood circulation (≈ 0.01%)*.

Of note, in a recent rabbit study, the development of a thermosensitive biodegradable slow-release drug delivery system (DDS) was reported. When placed subjconjunctivally, this device loaded with 2 mg infliximab reduced 90 day axon loss after alkali burn to the cornea from about 50% (n=4) to about 8% (n=3) (Zhou et al., 2021)—thus very similar to what is reported in this paper.

## Discussion

The inflammatory cytokine TNF-α seems to be playing a major role in the triggering of optic neuropathy after acute traumatic events in the eye (Roh et al., 2012; Cade et al., 2014). Most likely other inflammatory cytokines are involved as well, especially IL-6 and IL-1β (Paschalis et al., 2017), but the role of TNF-α in glaucoma has been under strong suspicion for over two decades (Madigan et al., 1996; Yuan et al., 2000). The development of monoclonal antibodies has helped to pinpoint the role of this cytokine in the process. Thus, it has been previously shown in animals that ganglion cell death after short exposure to high intraocular pressure can be blocked by etanercept, a TNF-α antibody (Roh et al., 2012). In the likely IOP-independent, rapid inflammatory pathway described by us in animals, infliximab and adalimumab similarly show a high level of ganglion cell protection (Cade et al., 2014; Paschalis et al., 2017). Whether these seemingly separate pathophysiological pathways to optic nerve damage (IOP-independent and IOP-triggered) are truly separate or merge into one, remains to be determined.

When interpreting the results of past alkali burn or corneal surgery experiments, the timing of events is important. Thus, TUNEL staining showing retinal cell apoptosis 3 days after the corneal burn should reflect only initial, rapid, seemingly IOP-independent events, whereas the optic nerve degeneration after 50 days (in earlier experiments 90 days (Paschalis et al., 2017) might reflect not only this first phase but also the second, IOP-related phase during the healing process (increased outflow resistance, etc.). Under any circumstances, both phases might be blocked with prompt delivery of TNF-α inhibitors already on the market, and this therapeutic possibility deserves to be subjected to further evaluation. Here we have tried to evaluate the efficacy, toxicity and feasibility of subjconjunctival administration of these medications in a corneal alkali burn model.

The potential toxicity of TNF-α antibodies to the retina first received attention two decades ago, triggered by the introduction of the intravitreal route of administration to target wet macular degeneration. Thus, in one such rabbit study, *<u>intravitreal</u>* adalimumab did not appear toxic in concentrations of 0.5 or 5 mg (Tsilimbaris et al., 2009)—these results being pertinent here.

Infliximab was tested intravitreally in a similar rabbit study and was found to be non-toxic in doses up to 2.0 mg (Theodossiadis et al., 2009). Likewise, intravitreal infliximab was found to be safe in another rabbit study at a dose of up to 1.7 mg (Giansanti et al., 2008). However significant toxicity has been encountered in humans after intravitreal injection of infliximab with doses of 0.5–2.0 mg, which underscores the possible importance of interspecies differences (Yu et al., 2006; Rosenfeld & Goodman 2009; Giganti et al., 2010).

Not only biologics but also corticosteroids are neuroprotective against acute ganglion cell apoptosis and should be useful drugs in this respect. In our previous experiments in animals the short-acting dexamethasone, for unknown reasons, showed less efficacy than the more long-acting Kenalog^®^ preparation of triamcinolone, in the clinically most commonly used concentrations (Paschalis et al., 2017). However, the well-known complications of local corticosteroids, particularly their suppressive effect on corneal wound healing, limit their use post events to modest concentrations or short duration. The toxicity of steroids has also been previously studied in intravitreal rabbit experiments; 0.5 mg or 1.0 mg triamcinolone into the vitreous did not produce morphological changes in the retina but 4.0 mg, 8.0 mg, and 20 mg produced toxic effects in the outer retina (Yu et al., 2006). Another study on intravitreal triamcinolone confirmed that 4.0 mg was retinotoxic and suggested that the vehicle was to blame (Lang et al., 2007).

It is difficult to compare the results from these toxicology studies with the outcomes from our subconjunctival administrations, but they seem grossly compatible with our findings of a single injection of 4.0 mg adalimumab being well-tolerated subconjunctivally, while 40 mg is not. The computerized exercise on bioavailability (see under Results) supports the intuitive assumption that a lower dose of 4.0 mg adalimumab to the eye would have less systemic toxicity than a subcutaneous injection of 40.0 mg of the same drug. Moreover, subconjunctival administration of adalimumab achieves slower drug release into the retina and improves drug bioavailability.

This could potentially reduce the need for repeated injections, as presently required for intravitreal administration of other inhibitors.

Not only severe trauma but also less invasive standard corneal surgery (PK, KPro, laceration repair, etc.) can lead to some rapid upregulation of TNF-α in the eye and ganglion cell apoptosis, which may, at least in part, contribute to subsequent secondary glaucoma (Chen et al., 2020).

Even though such routine surgery rarely triggers as much inflammation as severe trauma, it may make sense to apply similar prophylactic principles if the patients are suspected to be at risk.

The clinical implications of our combined clinical experience with past and more recent animal experiments point to the need for heavier and more sustained anti-inflammatory medication than is presently utilized after acute ocular trauma or routine surgery. *<u>Emphasis should be on very</u> <u>prompt delivery after the acute event</u>*, with administration of the drug to an easily accessible part of the eye where it can still reliably protect the retinal ganglion cells—pointing to subconjunctival application as practical and relatively safe. A more detailed pharmacokinetic study on the relative efficacy of various routes of administration of TNF-α inhibitors was beyond the scope of this investigation, however, one may cautiously draw some analogies from work on bevacizumab, the full-length monoclonal antibody against vascular endothelial growth factor (VEGF). Intravitreal injection was by far the most effective route of administration for retinal effect but subconjunctival injection was sufficient and is less dangerous. Both administration pathways resulted in similar systemic exposure, with the subconjunctival route better targeting the cornea (Nomoto et al., 2009). Thus, a subconjunctival injection of anti-TNF-α drugs would be safer and more practical than intravitreal injection for a corneal surgeon and would also be effective against corneal complications.

There were several limitations in this study. For example, the alkali burns in rabbits resulted in a lower percentage of affected ganglion cells than in a mouse model used in previous studies (Paschalis et al., 2017; Zhou et al., 2017). This seems to be attributable to ocular size differences between models. However, at 3 months, the cumulative damage to the retina and optic nerve in rabbits was significant, which suggests that the rabbit model of alkali burn is still relevant in assessing neuroprotective therapies. Also, pilot experiments to determine dose toxicity were not performed in triplicate, however, all critical data, especially on the protective effect of 4 mg adalimumab, were obtained in triplicate. In addition, although the method of obtaining IOP is not commercially available, as it was developed in-house and hence was not validated by any outside authority, we performed all necessary tests to ensure the validity of IOP readings using hydrostatic pressure columns. Moreover, the sensor used in this system is commercially available and validated. We have described results involving the use of this sensor in several previously published papers and in our experience, it has provided more reliable IOP data in injured corneas than any other conventional indirect tonometer. Another limitation was that the loss of retinal ganglion cells was not specifically visualized using Brn3a antibody. Instead, we performed H&E and PPD staining to capture the cumulative damage to the retinal and optic nerve. In this study, rabbit retinas were analyzed using cryosections and not flat mount imaging. Although flat mount retinal analysis would be ideal, rabbit eyes are large and very challenging to process for flat mount analysis. To overcome this limitation, we quantified the cumulative damage to the optic nerve axons using PPD. However, according to previous publications RGC loss precedes axonal loss and hence, our data may underrepresent the true loss of RGCs. In addition, we did not perform cryopreservation of the retinal tissue for cryosectioning, which may have caused tissue aberrations or abnormalities. Cryopreservation is typically performed by our laboratory in mouse eyes, but previous attempts to cryopreserve rabbit eyes have led to collapse of the globe and retinal detachment. To avoid this issue, cryopreservation was not attempted in this study. Perhaps a modified protocol of cryopreservation of rabbit eyes may help avoid this complication in the future. Finally, ERG and OCT assessments were not performed in chemically burned eyes due to lack of corneal clarity. Instead, retinas were analyzed histologically post-mortem.

On the positive side, the low level of toxicity of adalimumab corroborates previous findings (Tsilimbaris et al., 2009). Also, the substantial protective effect on the retinal cells by the biologics, in various concentrations, is very high and very persuasive (**Fig. 4).** Thus, particularly 4 mg adalimumab subconjunctivally seems to suppress apoptosis almost completely (and be locally non-toxic). Particularly encouraging are the more final 3-month results on both ganglion cells and nerve axons. Based on the results of these and earlier animal experiments, it may be reasonable to initiate human studies with biologic, low-risk anti-inflammatory regimens following corneal surgery or unexpected trauma. In support, we have already shown in patients that infliximab can have a dramatic effect in preventing corneal tissue melt around a keratoprothesis (Dohlman 2022). Also, the FDA has approved the successful use of subcutaneous adalimumab in non-infectious uveitis. A pilot clinical study on the protective effects on the eye should be the next urgent logical step. It should be noted, however, that the role of the intraocular pressure as a triggering factor in <u>open-angle glaucoma</u> is not questioned here, where we might also encounter a separate, by-definition mechanism, for the secondary glaucomatous picture resulting from a single, acute, often-violent event acting as a triggering factor. Lastly, the large protein-based anti-inflammatory should have vast advantages when compared to the low weight, inexpensive anti-inflammatories (namely triamcinalone, dexamethasone, and prednisone) in terms of their monoclonal qualities. Due to their precision in identifying the inflammatory cytokine along with a broad spectrum of activities, they will remain more expensive but valuable for their specificity. The standard broad-spectrum anti-inflammatories will still be widely used, although they are well-known for pressure rising, cataract formation, and osteoporosis. Meanwhile, the biologics are more difficult to manufacture and have a propensity for making complexes, which can make them susceptible to immunogenicity. Nevertheless, they will continue to revolutionize the field for a long time and both classes will undoubtedly have to co-exist for a long time.

### Significance statement

This study demonstrates that low dose (4 mg) subconjunctival administration of the TNF-α antibody adalimumab can be safe and very effective in protecting against retinal damage in a rabbit model. This suggests the possibility of using this inhibitor as prophylaxis against late secondary glaucoma, with minimal systemic risk for the patient.

## Acknowledgements

The authors thank Leonard Levin, MD, PhD, and Anders Heijl, MD, PhD for valuable advice.

---

**Table 1.** Summary of results of testing adalimumab, infliximab and triamcinolone, injected subconjunctivally, for **(A)** toxicity and **(B)** efficacy in reducing inflammation of the retinal cells and optic nerve degeneration, after alkali burn to the corneas of rabbits. (The results from Zhou, Robert et al. drug delivery systems with infliximab are added for comparison. See Refs 65, 75). Adalim=adalimumab, Inflix=infliximab, Triam=triamcinolone, DDS=drug delivery system (inserted subconjunctivally)

**Toxicity to retina or optic nerve from subconjunctival injection of drugs in normal rabbit eyes**
| **Drug concentration** | **Observation days/Toxicity** |  | **Summarized from** |
| --- | --- | --- | --- |
| Adalim 0.4 mg | 50 | - | Fig 1 |
| Adalim 4 mg | 50 | - | Figs 1, 2 |
| Adalim 40 mg | 50 | ++ | Figs 1, 2 |
| Inflix 1 mg | 50 | - | Fig 1 |
| Inflix 10 mg | 50 | - | Fig 1 |
| Inflix 100 mg | 50 | - | Figs 1, 2 |
| Saline (sham) | 50 | - | Figs 1, 2 |

**Efficacy: Protective effect on retina and optic nerve after burn to cornea**
| **Drug concentration** | **Observation days/Efficacy** |  |  |  | **Summarized from** |
| --- | --- | --- | --- | --- | --- |
| Adalim 0.4 mg | 3 |  |  |  |  |
| Adalim 4 mg | 3 | +++ | 90 | +++ | Figs 3, 4 |
| Adalim 40 mg | 3 |  |  |  |  |
| Inflix 1 mg | 3 |  |  |  |  |
| Inflix 10 mg | 3 |  |  |  |  |
| Inflix 100 mg | 3 | ++ |  |  | Fig 3 |
| Triam 20 mg | 3 | ++ |  |  | Fig 3 |
| Saline (sham) | 3 | - | 90 | - | Figs 3, 4 |
| (DDS' Inflix |  |  | 90 | +++ | p. 14, refs 65, 75) |

Supplemental Figure 1. Toxicity: Dark-adapted ERG assessment after subconjunctival injection of adalimumab and infliximab.Dark-adapted electroretinography (ERG) using 0.01, 3 and 10 cd.s/m^2^ light intensities after subconjunctival injection of **(A - C)** 40 mg adalimumab and **(D - F)** 100 mg infliximab. The contralateral eyes were used as internal controls, while saline (sham) injected eyes served as treatment controls. ERG quantification in 40 mg adalimumab injected eyes **(G)** Amplitudes **(H)** Implicit times. 100 mg infliximab injected eyes **(I)** Amplitudes and **(J)** Implicit times.Measurements were performed at baseline (7 days before injection) and 3, 7, 28, and 45 days after injection. **(G - J)** Quantification was performed for “a” and “b”-wave “A”mplitude and“T”ime responses at“0.01”, “3”, and “10” cd.s/m^2^ light intensities in injected eyes. High doses of subconjunctival injection of adalimumab and infliximab did not cause appreciable changes in the dark-adapted ERG responses, although optic nerve degeneration was evident at 40 mg of adalimumab with PPD staining. One animal per dose with serial measurements, total n=8.

Supplemental Figure 2. Toxicity: Light-adapted ERG assessment after subconjunctival injection of adalimumab and infliximab. Light-adapted electroretinography (ERG) using 3 cd.s/m^2^ flash and flicker light stimulation after subconjunctival injection of **(A, B)** 40 mg adalimumab and **(C, D)** 100 mg infliximab. (E,F) Amplitudes **and** corresponding implicit times of ERG in 40 mg adalimumab and (G,H) 100 mg infliximab injected eyes.Measurements were performed at baseline (7 days before injection) and 3, 7, 28, and 45 days after injection. **(E - H)** Quantification was performed for “a” and “b”-wave “A” amplitude and “T” Time responses at“ 3” cd.s/m^2^ “flash” and “flicker” light intensity in injected eyes. High dose subconjunctival injection of adalimumab and infliximab did not cause appreciable changes in the light-adapted ERG responses. One animal per dose with serial measurements, total n=8.

Supplemental Figure 3. Magnified optical coherent tomography of the retina 50 days after subconjunctival administration of adalimumab 40 mg.Representative high magnification *in vivo* optical coherence tomography (OCT) of the superior retinal quadrant at baseline and 50 days post subconjunctival injection with 40 mg adalimumab showing thinning of the neuroretina.

