## Supplementary material for "The prophylactic value of TNF-α inhibitors against retinal cell apoptosis and optic nerve axon loss after corneal surgery or trauma": Sup Figure 1

**Sup. Fig. 3. Representative OCT images of the superior retina before and after subconjunctival injection of 40mg adalimumab**

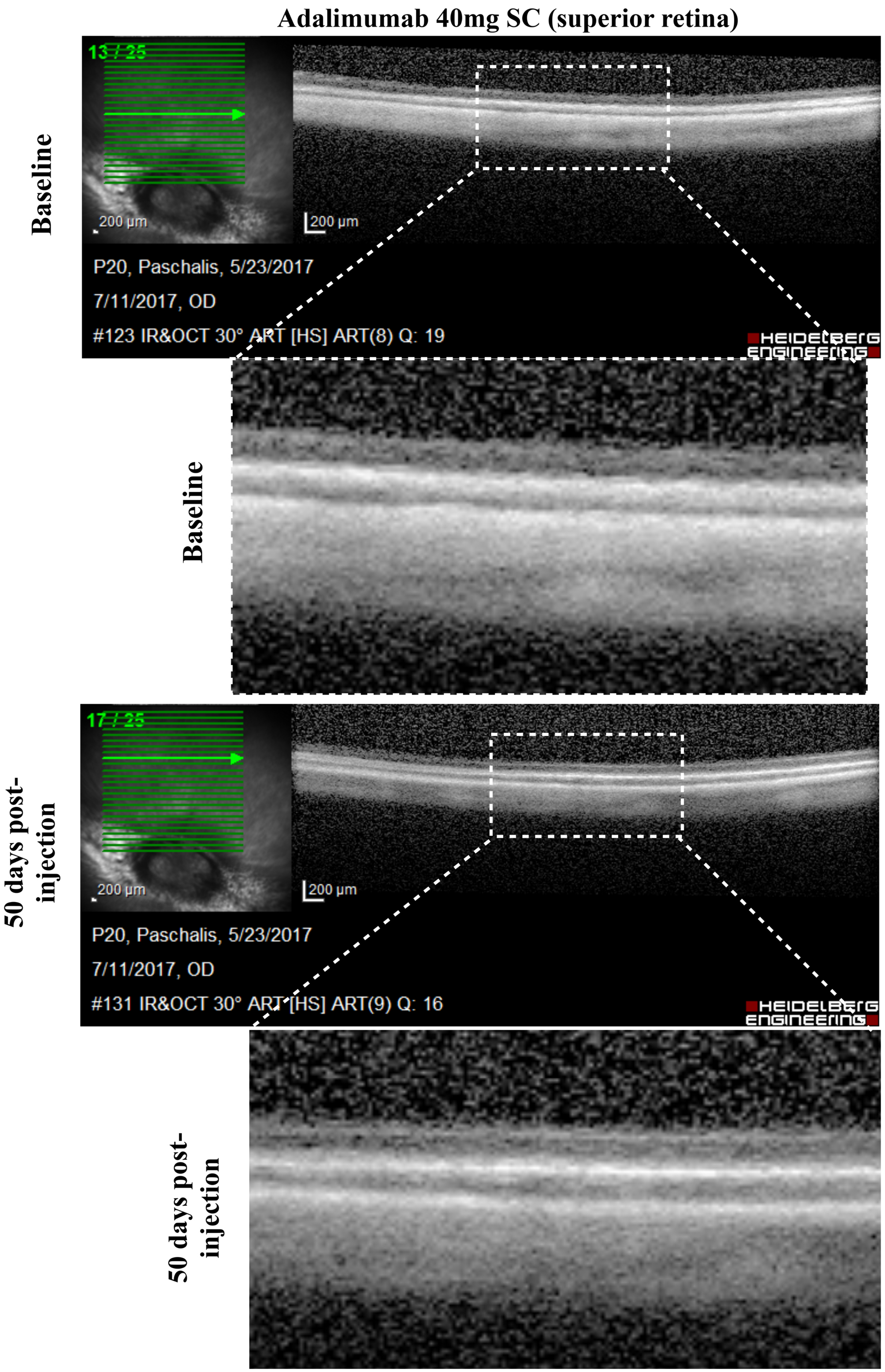
