## Supplementary material for "The prophylactic value of TNF-α inhibitors against retinal cell apoptosis and optic nerve axon loss after corneal surgery or trauma": Sup Figure 3

**Sup. Fig 2. Toxicity evaluation of subconjunctival injection of adalimumab or infliximab at maximum dose with light-adapted ERG**

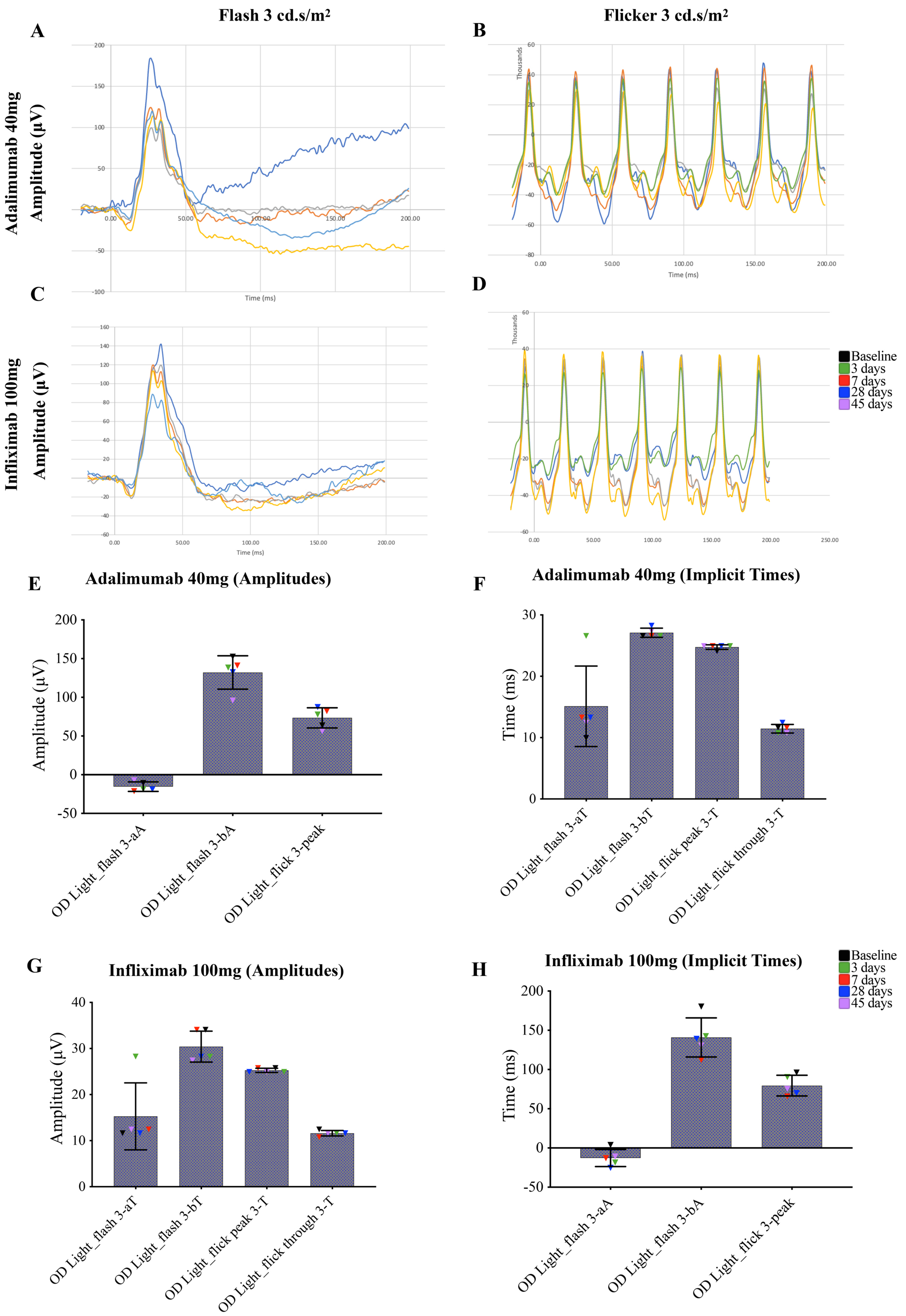
